# Association of GLOD4 with Alzheimer’s Disease in Humans and Mice

**DOI:** 10.1101/2024.06.07.597934

**Authors:** Olga Utyro, Olga Włoczkowska-Łapińska, Hieronim Jakubowski

**Author notes:** These authors contributed equally to this work (Olga Utyro, Olga Włoczkowska-Łapińska). Correspondence to: Hieronim Jakubowski, Department of Microbiology, Biochemistry and Molecular Genetics, Rutgers-New Jersey Medical School, International Center for Public Health, 225 Warren Street, Newark, New Jersey 07103, USA.

## Abstract

**Background:** Glyoxalase domain containing protein 4 (GLOD4), a protein of an unknown function, is associated with Alzheimer’s disease (AD). Three GLOD4 isoforms are known. The mechanism underlying GLOD4’s association with AD was unknown.

**Objective:** To assess GLOD4’s role in the central nervous system by studying GLOD4 isoforms expression in human frontal cerebral cortical tissues from AD patients and in brains of *Blmh*^−/−^5xFAD mouse AD model of AD.

**Methods:** GLOD4 protein and mRNA were quantified in human and mouse brains by western blotting and RT-qPCR, respectively. Mouse brain amyloid β (Aβ) was quantified by western blotting. Behavioral assessments of mice were performed by cognitive/neuromotor testing. *Glod4* gene in mouse neuroblastoma N2a-APPswe cells was silenced by RNA interference and Glod4 protein/mRNA, Aβ precursor protein (Aβpp)/mRNA, *Atg5*, *p62*, and *Lc3* mRNAs were quantified.

**Results:** *GLOD4* mRNA and protein isoforms were downregulated in cortical tissues from AD patients compared to non-AD controls. *Glod4* mRNA was downregulated in brains of *Blmh*^−/−^5xFAD mice compared to *Blmh*^+/+^5xFAD sibling controls, but not in *Blmh*^−/−^ mice without the 5xFAD transgene compared to *Blmh*^+/+^ sibling controls. The 5xFAD transgene downregulated Glod4 mRNA in *Blmh*^−/−^ mice of both sexes and in *Blmh*^+/+^ males but not females. Attenuated Glod4 was associated with elevated Aβ and worsened memory/sensorimotor performance in *Blmh*^−/−^5xFAD mice. Glod4 depletion in N2a-APPswe cells upregulated AβPP and downregulated autophagy-related *Atg5*, *p62*, and *Lc3* genes.

**Conclusions:** These findings suggest that GLOD4 interacts with AβPP and the autophagy pathway, and that disruption of these interactions leads to Aβ accumulation and cognitive/neurosensory deficits.

## INTRODUCTION

Alzheimer’s disease (AD) is the primary cause of dementia, affects 55.0 million individuals worldwide and ranks as the fifth leading cause of death globally [1]. In the United States alone, an estimated 6.9 million individuals aged 65 and older (2% of the US population) live with AD dementia today, and this number is expected to grow to 13.8 million by 2060. AD is characterized by the extracellular accumulation of amyloid β (Aβ) and the intracellular accumulation of neurofibrillary tangles of hyperphosphorylated tau protein leading to neuronal death. Mutations in amyloid β protein precursor protein (AβPP), presenilin 1 (PSEN1), and presenilin 2 (PSEN2) lead to the familial early-onset AD, which is relatively rare [2]. Although lifestyle and environmental factors have been recognized as modifiers of the susceptibility to AD [3], the causes of the most prevalent sporadic late-onset AD are largely unknown. Because there is no proven way to prevent AD and there is currently no cure available, it is important to identify factors influencing AD and their mode of action.

There is evidence suggesting that the glyoxalase domain containing protein 4 (GLOD4), a protein of unknown function, may play a significant role in the central nervous system (CNS) and may be linked to AD. For instance, an intronic SNP in *GLOD4*, rs2750012, was associated with increased risk of AD in the Arab population of northern Israel. This association has been replicated in metanalysis performed in seven independent GWAS datasets [4]. Levels of one of the Glod4 protein isoforms, isoform 3, were significantly elevated in APPSwDI/*NOS2^−/−^* mouse model of AD compared to controls [5]. Proteomic studies of microglial cells BV-2 showed that treatments with the anti-inflammatory drug polyphenol pentagalloyl glucose (PGG) downregulated GLOD4 while treatments with proinflammatory LPS/IFNγ upregulated GLOD4 [6]. In addition to GLOD4, PGG downregulated septin-7, ataxin-2, and adenylosuccinate synthetase isozyme 2 (ADSS), which were linked in earlier studies to neurodegenerative diseases such as AD, Parkinson’s, Huntington’s, Down’s syndrome, and frontotemporal dementia [6]. *GLOD4* was also identified among 38 duplicated genes in multiple cases of autism spectrum disorder (ASD) [7].

The *GLOD4* gene (C17orf25) is found in a region on human chromosome 17 showing high heterozygosity in human hepatocellular carcinoma [8]. Human GLOD4 is expressed in most tissues, including the brain [8] (https://www.proteinatlas.org/ENSG00000167699-GLOD4). GLOD4 belongs to the glyoxalase gene family that includes glyoxalase 1 (GLO1), which detoxifies methylglyoxal [9]. Methylglyoxal, a byproduct of glucose metabolism [10] responsible for protein glycation can causes misfolding [11]. Glycated proteins easily aggregate, which can give rise to amyloid plaques, a hallmark of AD [12]. Although human GLOD4 was suggested to take part in the methylglyoxal detoxification [5], this still is to be shown. A yeast two-hybrid screen showed that GLOD4 protein interacts with mitochondrial ADP-ribose pyrophosphatase NUDT9, which may regulate cell growth [13].

The human GLOD4 protein has 3 isoforms, differing in amino acid sequence (www.uniprot.org/blast/). The isoform 1 consists of 313 amino acids and contains motifs of the glyoxalase domain [9], antibiotic resistance domain, and a domain characteristic of the family of proteins resistant to the cytotoxic anticancer drug bleomycin (www.ncbi.nlm.nih.gov/protein/NP_057164.3).

Bleomycin hydrolase (BLMH) plays a key role in the CNS and is linked to AD. For example, BLMH can process AβPP to Aβ [14] and to further process Aβ [15]. Enzymatic activities of BLMH are significantly downregulated in the human AD brain [16]. *Blmh*^−/−^ mice show astrogliosis, behavioral changes [17], and diminished ability to detoxify Hcy-thiolactone, which elevates brain Hcy-thiolactone level and increases its neurotoxicity [4]. Proteomic studies of *Blmh*^−/−^ mouse brain show that Blmh interacts with diverse cellular processes, such as synaptic plasticity, cytoskeleton dynamics, cell cycle, energy metabolism, and antioxidant defenses that are essential for brain homeostasis [18]. *Blmh*^−/−^5xFAD mouse model of AD shows exacerbated cognitive/neuromotor deficits [19, 20]. These neurological impairments resulted from the disruption of interactions of Blmh with AβPP and the Phf8/H4K20me1/mTOR/autophagy pathway, which lead to Aβ accumulation and cognitive/neuromotor deficits [20].

Here we examined the role of GLOD4 in the CNS by studying GLOD4 expression in AD brains. We also examined Glod4 expression in *Blmh*^−/−^5xFAD mouse model of AD in relation to cognitive/neuromotor performance in these mice and biochemical consequences of *Glod4* gene silencing in mouse neuroblastoma N2a-APPswe cells. Our findings show that GLOD4 is important for CNS homeostasis.

## MATERIALS AND METHODS

### Human brain samples

Human frontal cerebral cortical tissues were obtained at autopsy by the Case Western Reserve University Brain Bank under an IRB-approved protocol, from clinically and pathologically confirmed cases of AD (n = 6) using criteria set up by the National Institute of Aging (NIA) and Consortium to Establish a Registry for Alzheimer’s Disease (CERAD) [21]. Brain tissues were also obtained from non-AD control patients (n = 6). Samples of frontal cortex, coded prior to shipment to keep laboratory personnel blinded as to diagnosis, were shipped on dry ice to the New Jersey Medical School and stored at−80◦C until used. The AD patients were 84±8-year-old, 67% female, post-mortem interval was 8±5 h. Controls were 77±6-year-old (*P* = 0.100), 83% female (*P* = 0.549), post-mortem interval was 16±8 (*P* = 0.103).

### Mice

*Blmh*^−/−^5xFAD mice, obtained by crossing *Blmh*^−*/*−^ [22] and 5xFAD [23] animals, were characterized in an earlier study [20]. *Blmh* and 5xFAD genotypes were established by PCR using primers listed in **Table 1**. Mice were kept at the Rutgers-New Jersey Medical School Animal Facility as previously described [20]. 5xFAD mice carry a transgene with the K670 N/M671 L (Swedish), I716V (Florida), and V717I (London) mutations in human AβPP(695), and M146 L and L286V mutations in human PS1 associated with familial early-onset AD. 5xFAD mice accumulate elevated levels of Aβ42 beginning around 2 months of age [23]. Four groups of 5-month-old mice of both sexes were used in experiments: *Blmh*^−*/*−^, their wild type *Blmh*^+*/*+^ littermates, *Blmh*^−*/*−^5xFAD, and their *Blmh*^+*/*+^5xFAD littermates. The groups were derived from multiple litters and equal number of females and males were used in each group. Animal procedures were approved by the Institutional Animal Care and Use Committee at Rutgers-New Jersey Medical School.

**Table 1.**
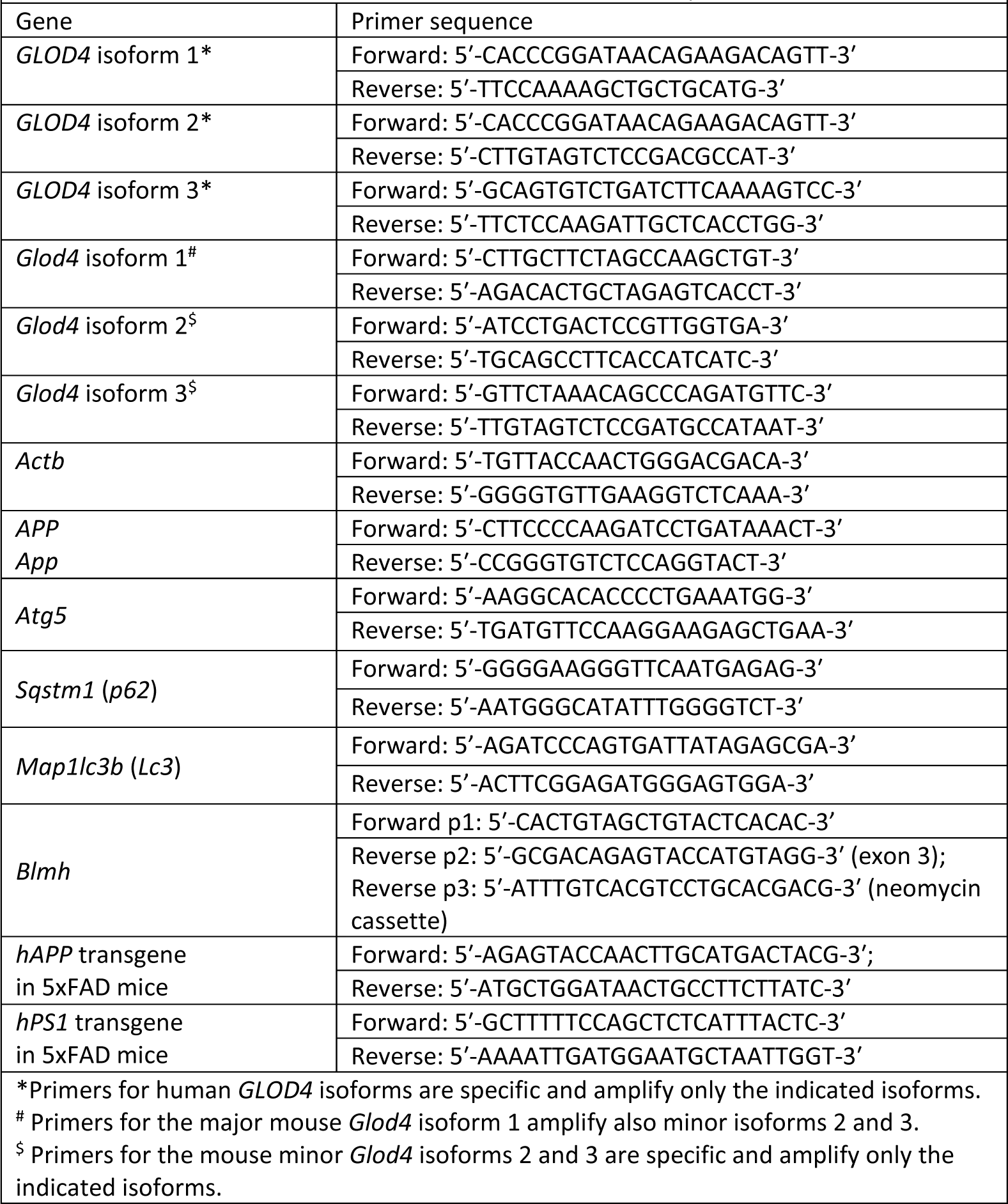
Primers used for PCR or RT-qPCR.

### Behavioral testing

Cognitive and neuromotor performance was evaluated using the novel object recognition (NOR) [24] and the hindlimb [25] tests as previously described [20]. Briefly, NOR evaluates recognition memory in two sessions. During the familiarization session, a mouse freely explores two identical objects for 20 s. Six hours later, during the test session with one object replaced by a novel object, a mouse is allowed to explore for 20 s. The time spent exploring each object is calculated from video recordings. Cognitively healthy mice spend more time exploring a novel object, while cognitively impaired mice do not differentiate between novel and familiar objects.

The hindlimb test evaluates neurodegeneration. A mouse is suspended by tail and videorecorded for 10 s in three trials over three days. Hindlimb clasping is scored from 0 to 3: score 0, hindlimbs splayed outward and away from the abdomen; score 1, one hindlimb retracted inwards towards the abdomen for at least 50% of the observation period; score 2, both hindlimbs partially retracted inwards towards the abdomen for at least 50% of the observation period; score 3, both hindlimbs completely retracted inwards towards the abdomen for at least 50% of the observation period. The scores were averaged for the three trials. A higher score indicates worse neurodegeneration.

### Cell culture and Glod4 siRNA transfection

Mouse neuroblastoma N2a-APPswe cells, harboring a human AβPP transgene with the K670N and M671L Swedish mutations associated with familial early-onset AD [26] were grown (37°C, 5% CO_2_) in DMEM/F12 medium (Thermo Scientific) supplemented with 5% FBS, non-essential amino acids, and antibiotics (penicillin/streptomycin) (MilliporeSigma).

Cell monolayers (50-60% confluency), washed twice with PBS, were transfected with *Glod4*-targeting siRNAs (siRNA Glod4 #1: Cat. #43390816 ID s84641; siRNA Glod4 #2: Cat. #4390816 ID s84642, Thermo Scientific); siRNA scrambled (siRNAscr; Silencer R Negative Control siRNA #1, AM4635, Ambion) in Opti-MEM medium with Lipofectamine RNAiMax (Thermo Scientific) for 48 hours.

### Western blots

Proteins from human cerebral cortex and whole mouse brain were extracted using RIPA buffer (MilliporeSigma) with Protease Inhibitor cocktail (Sigma P8340) as previously described [20]. Proteins were separated by SDS-PAGE on 16% acrylamide gels immobilized by transfer to a PVDF membrane (Cat. #1620177, Bio-Rad) using a Trans-Blot Turbo Transfer System (Trans-Blot ®Turbo^TM^, Bio-Rad) according to manufacturer’s specifications (20 min, 120 mA, 25 V), and quantified with the following antibodies: anti-GLOD4 (Novus Biologicals NBP2-60713, 1:1,000 dil.), anti-GAPDH (MilliporeSigma PLA0125, 1:4,000 dil.), anti-Glod4 (Abcam AB188371, 1:1,000 dil.), anti-App (Abcam ab126732, 1:1,000 dil.), anti-β-actin (Abcam ab8227, 1:3,500 dil.), or anti-Gapdh (CST #5174, 1:2,000) for 1 hour. Membranes were washed three times with TBST, 10 min each, and incubated with goat anti-rabbit IgG secondary antibody conjugated with horseradish peroxidase (CST, #7074S, 1:3,000). Positive signals were detected using Western Bright Quantum-Advansta K12042-D20 and GeneGnome XRQ NPC chemiluminescence detection system. Bands intensity was calculated using Gene Tools program from Syngene.

### RNA isolation, cDNA synthesis, RT-qPCR analysis

Total RNA was isolated using Trizol reagent (MilliporeSigma). RNA concentration was measured using NanoDrop (Thermo Fisher Scientific).and integrity was assessed by agarose gel (1%) electrophoresis. cDNA was synthesized using Revert Aid First cDNA Synthesis Kit (Thermo Fisher Scientific) according to manufacturer’s protocol. RT-qPCR reactions (2 ng/µL cDNA, primers 0.3 µM each, iTaq 5 µL in a total volume of 10 µL) were performed in two technical replicates using SYBR Green Mix and the CFX96 Touch Real-Time PCR Detection System (Bio-Rad). RT-qPCR primer sequences are listed in **Table 1**.

For mouse *Actb*, *Gapdh*, *Glod4_1*, *Glod4_2*, *Glod4_3* the following reaction conditions were used: (1) initial denaturation 2 min, 95°C; (2) denaturation 15 s, 95°C; (3) primer annealing (temperature (T)), 15 s; (4) primer extension 10 s, 72°C; melting curve 65-95°C. T = 60°C for *Glod4_1*, *Glod4_2*, *Glod4_3*, *App*, *Atg5*, *p62*; T = 56°C for *Actb*. The cycle of denaturation, annealing, and extension was repeated 40 times.

For human *GAPDH*, *GLOD4_1*, *GLOD4_2*, *GLOD4_3* the following reaction conditions were used: initial denaturation 2 min, 95°C; denaturation 35 s, 95°C; primer annealing 10 s, 62°C; primer extension, 30 s, 72°C; melting curve 65-95°C. The cycle of denaturation, annealing, and extension was repeated 40 times.

The 2^(-ΔΔCt)^ method was used to calculate the relative expression levels [27]. Data were analyzed using CFX Manager™ Software and Microsoft Excel.

### Data analysis

Normality of distributions was evaluated with the Shapiro-Wilk’s statistic. For variables with normal distributions, the results were calculated as mean±SD for variables with normal distributions or medians for non-normally distributed variables. A parametric unpaired t test was used for comparisons between two groups of variables with normal distribution. A Mann-Whitney rank sum test was used for comparisons between two groups of non-normally distributed variables. Data were analyzed using GraphPad Prism7 software (GraphPad Holdings LLC, San Diego CA, USA, https://www.graphpad.com) and MS Excell.

## RESULTS

### GLOD4 is downregulated in AD brains

Three isoforms of GLOD4 protein exist in humans: 34.8 kDa isoform 1 (GLOD4_1), 33.2 kDa isoform 2 (GLOD4<u>_</u>2), and 21.3 kDa isoform 3 (GLOD4<u>_</u>3). We quantified mRNAs for *GLOD4* isoforms in frontal cerebral cortical tissues from brains of six AD patients and six non-AD controls by using RT-qPCR. We found that *GLOD4_1* mRNA level was the highest in control human brains; *GLOD4_2* and *GLOD4_3* mRNAs were present at 10-fold lower levels (**Figure 1A**). Similar pattern of *GLOD4* mRNA isoforms expression was seen in AD brains (**Figure 1A**). However, relative levels of *GLOD4* mRNA isoforms were significantly reduced in AD brains: from 10.31 to 3.83 (by 63%) for GLOD4_1 mRNA, from 1.00 to 0.51 (by 49%) for *GLOD4_2* mRNA, and from 1.10 to 0.30 (by 73%) for *GLOD4_3* mRNA (**Figure 1A**).

**Figure 1.**
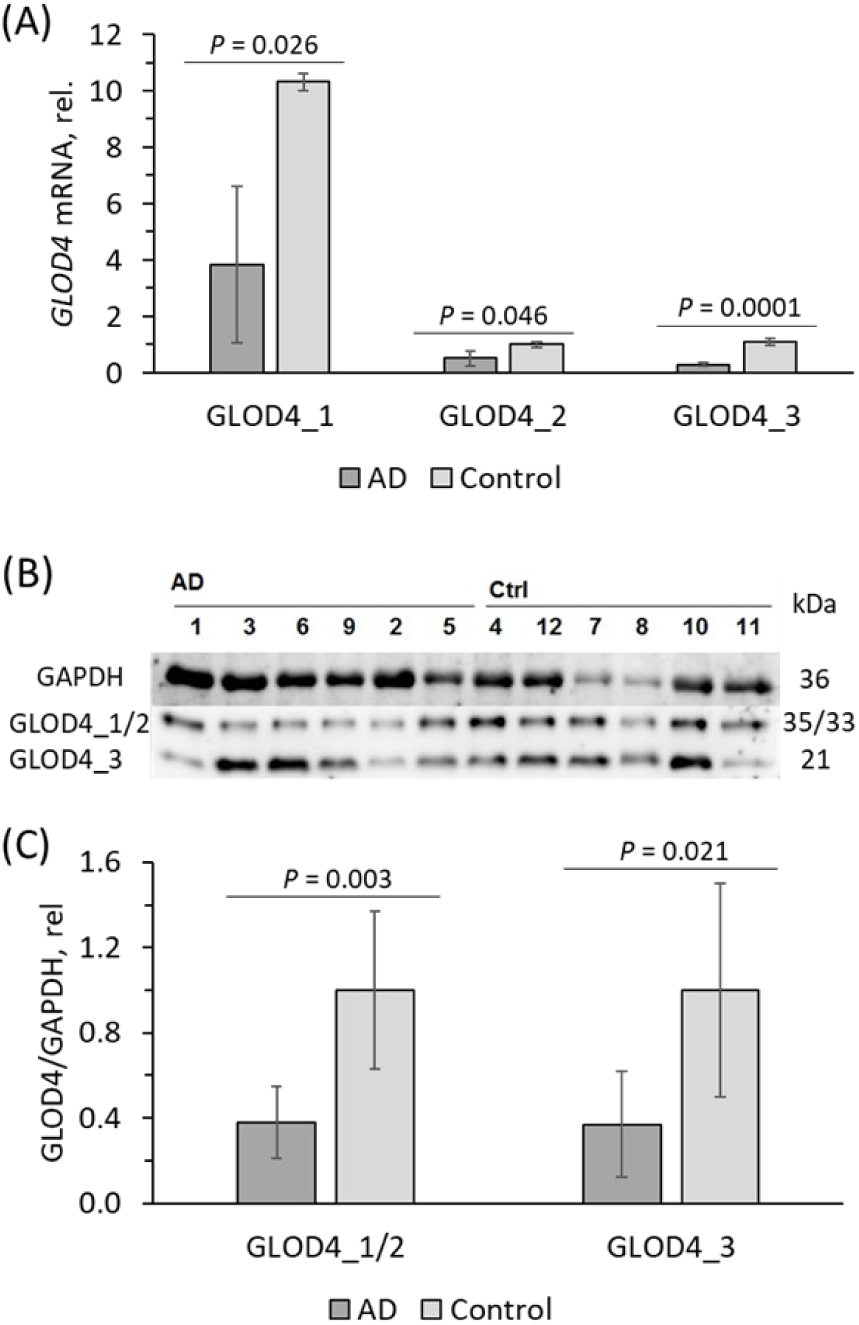
Reduced expression of GLOD4 isoforms in brains of Alzheimer’s disease patients. Isoforms of *GLOD4* mRNA and protein were quantified in the frontal cortex samples from AD patients (n = 6) and non-AD controls (n = 6). (A) Bar graph shows *GLOD4* mRNA quantification by RT-qPCR using *ACTB* mRNA as a reference. (B, C) GLOD4 protein isoforms were quantified by western blotting using GAPDH protein as a reference. (B) Representative image of a western blot. (C) Bar graph shows GLOD4 protein quantification. The experiment was performed twice. *p*-values were calculated by unpaired two-sided Student t test.

GLOD4 protein, quantified by western blotting, was similarly downregulated in frontal cerebral cortex tissues from human AD brains. SDS-PAGE analyses revealed two bands reacting with anti-GLOD4 antibody in the cortical brain protein extracts: an upper band of 35/33 kDa corresponding to GLOD4_1 and GLOD4_2 and a lower band of 21 kDa corresponding to GLOD4_3, based on the molecular weight (**Figure 1B**). Quantification showed that GLOD4_1/2 protein level was significantly reduced in AD brains compared to controls (0.62 vs. 1.00) as was the level of GLOD4_3 protein (0.63 vs. 1.00) (**Figure 1C**).

### Glod4 level is associated with Aβ and cognitive/neuromotor performance in Blmh^−/−^5xFAD mice

To determine whether downregulated GLOD4 levels seen in AD patients are associated with hallmarks of AD, we quantified *Glod4* mRNA level in *Blmh^−/−^*5xFAD mice, a model of human AD; *Blmh*^+/+^5xFAD littermates were used as controls [20]. The choice of the *Blmh^−/−^5xFAD* mouse model [20] for studies of *Glod4* was based on our previous findings showing that Blmh activity was reduced in human AD brains [16]. Because Aβ starts to accumulate in 5xFAD mice at 2 months [23], this model allowed examination of a relationship between Glod4 expression, behavioral performance, and Aβ accumulation at an early age. Earlier analyses of gene expression in mouse liver showed that *Glod4_1* mRNA is a major isoform while *Glod4_3* is a minor isoform while *Glod4_2* isoform was not detectable by PCR and that *Glod4_3* mRNA isoform was not expressed *Blmh*^−/−^ mouse liver [18].

Using RT-qPCR we found that *Glod4_1* was the most abundant isoform and *Glod4_3* was present at 25-fold lower level than *Glod4_1* in the mouse brain (0.041 vs. 1.00, Table 2). *The Glod4_2* mRNA isoform was essentially absent in the mouse brain, with an estimated level 1250-fold (*0.0008 vs. 1.00*) and 51-fold (*0.0008 vs. 0.041*) lower than those of *Glod4_1* and *Glod4_3* mRNAs, respectively (Table 2).

**Table 2.** Relative expression of Glod4 mRNA isoforms in the mouse brain.

| Table 2. Relative expression of *Glod4* mRNA isoforms in the mouse brain |  |  |  |  |  |
| --- | --- | --- | --- | --- | --- |
| Isoform | Relative expression * |  |  |  |  |
|  | *Blmh*<sup>+/+</sup> mice | *Blmh*<sup>-/-</sup> mice | Fold change | *P*<sub>*Blmh* genotype</sub> | *P*<sub>*Glod4* isoform</sub> |
| *Glod4_1* | 1.00±0.26 | 1.24±0.53 | 1.24 | 0.340 | 1.0 |
| *Glod4_2* | 0.0008<sup>#</sup> |  |  |  |  |
| *Glod4_3* | 0.041±0.014 | 0.00003±0.000006<sup>s</sup> | 1,255 | 0.004 | 1E.-08 |
| * Mouse brain *Glod4* mRNA isoforms were quantified by RT-qPCR using *Actb* mRNA as a reference and primers listed in **Table 1**. Six to seven-week-old *Blmh*<sup>+/+</sup> (n=12) and *Blmh*<sup>-/-</sup> (n = 10) mice of both sexes were used in the experiments. Signals for *Glod4_3* were observed in brain samples from all wild type *Blmh*<sup>+/+</sup> mice and in two out of ten *Blmh*<sup>-/-</sup> mice. <sup>#</sup> Estimated from RT-qPCR measurements in duplicates with 10 samples containing 5-80 ng cDNA prepared from wild type *Blmh*<sup>+/+</sup> mouse brain RNA. Signals for *Glod4_2* were observed for 4 out of 10 samples. |  |  |  |  |  |

We found that *Glod4_1/3* mRNA was significantly downregulated in brains of *Blmh^−/−^*5xFAD mice compared to *Blmh*^+/+^5xFAD sibling controls (**Figure 2A**), thus recapitulating the findings in the human AD brain (**Figure 1A**). Low Glod4 expression level in *Blmh^−/−^*5xFAD brains was accompanied by elevated Aβ level (**Figure 2B**).

**Figure 2.**
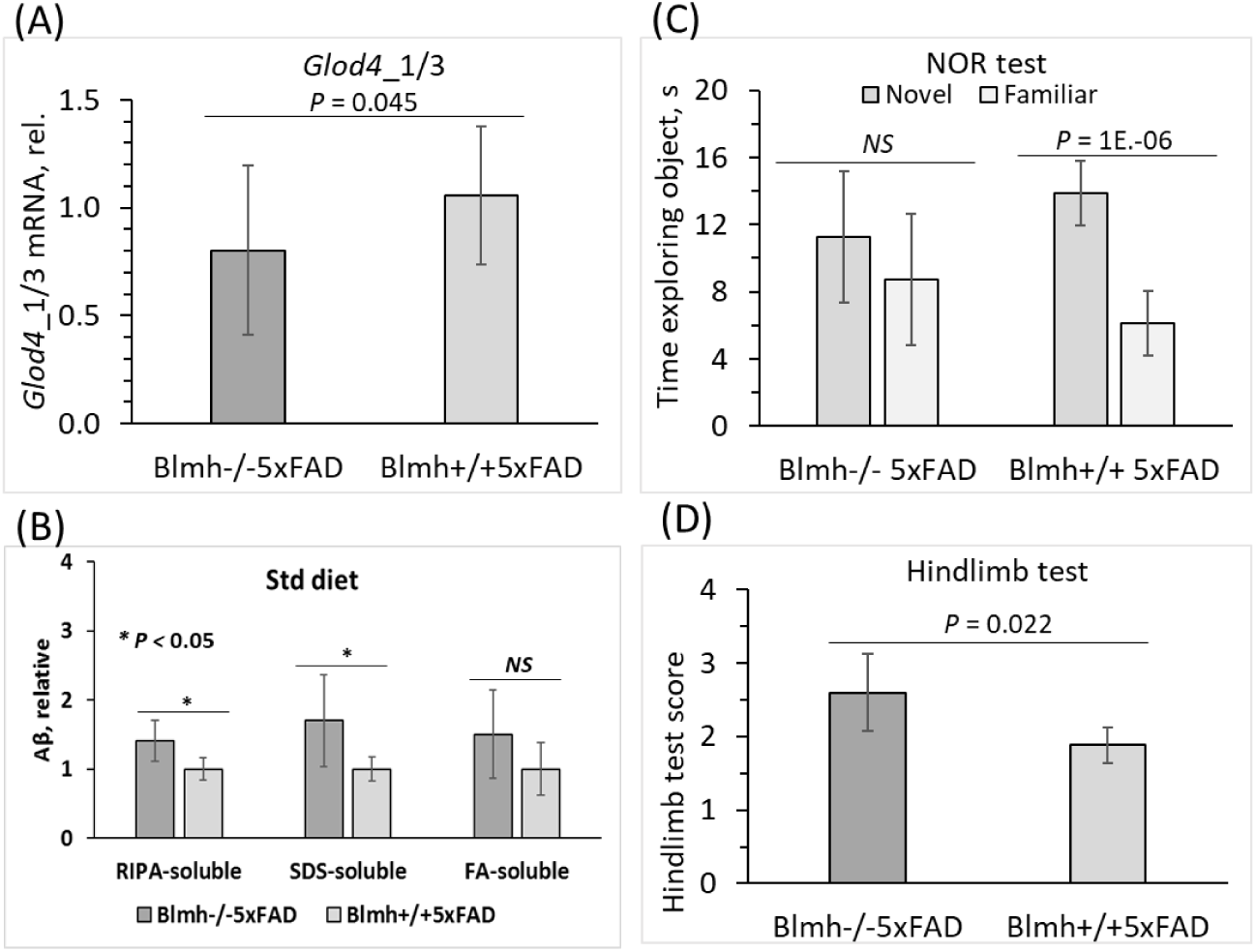
Reduced expression of brain *Glod4*_1/3 mRNA is associated with elevated Aβ and impaired cognitive/neuromotor performance in a mouse model of Alzheimer’s disease. Five-month-old mice of both sexes were used in the experiments: *Blmh*^−/−^5xFAD (n = 22) and their *Blmh*^+/+^5XFAD siblings (n = 15). Bar graphs illustrate quantification of (A) *Glod4*_1/3 mRNA using *Actb* mRNA as a reference, (B) Aβ, and cognitive/neuromotor performance in (C) NOR and (D) hindlimb tests. Panels B, C, and D were reproduced with permission from [20]. Data points for each mouse group represent mean ± SD of two to four independent measurements for each mouse. *p*-values for *Glod4*_1/3 mRNA and the hindlimb test were calculated by an unpaired two-sided Student t test. *p*-values for the NOR test were calculated by the paired two-sided Student t test. *p*-values for Aβ were calculated by one-way ANOVA with Tukey’s multiple comparisons test. *NS*, not significant.

Behavioral testing showed that *Blmh^−/−^*5xFAD mice with lower Glod4 level also had neurological impairments characteristic of human AD. Specifically, these mice spent the same time exploring novel and familiar objects, i.e., did not differentiate between novel and familiar objects in the NOR test, which shows impaired recognition memory. In contrast, *Blmh*^+/+^5xFAD mice that had higher Glod4_1 level spent more time exploring a novel object, i.e., had normal preference for novelty (**Figure 2C**).

In the hindlimb clasping test, *Blmh*^−/−^5xFAD mice that showed downregulated Glod4_1 level also showed significantly higher scores (indicating neuromotor deficits) compared to *Blmh*^+/+^5xFAD animals that had higher Glod4_1 (**Figure 1A**) and lower scores (i.e., better neuromotor performance) (**Figure 2D**).

### 5xFAD transgene affects Glod4 expression in Blmh^−/−^ and wild type Blmh^+/+^ mice

To assess whether the 5xFAD transgene affects *Glod4* expression, we quantified *Glod4* mRNA in *Blmh*^−/−^ and *Blmh*^+/+^ mice without the 5xFAD transgene. We found that *Glod4_1/3* mRNA was significantly downregulated in *Blmh*^−/−^5xFAD mice compared to non-transgenic *Blmh*^−/−^ animals (**Figure 3A**). We also found that the effect of the 5xFAD transgene on *Glod4* mRNA expression in *Blmh*^−/−^ mice was sex-independent, similar in females and males (**Figure 3B**).

**Figure 3.**
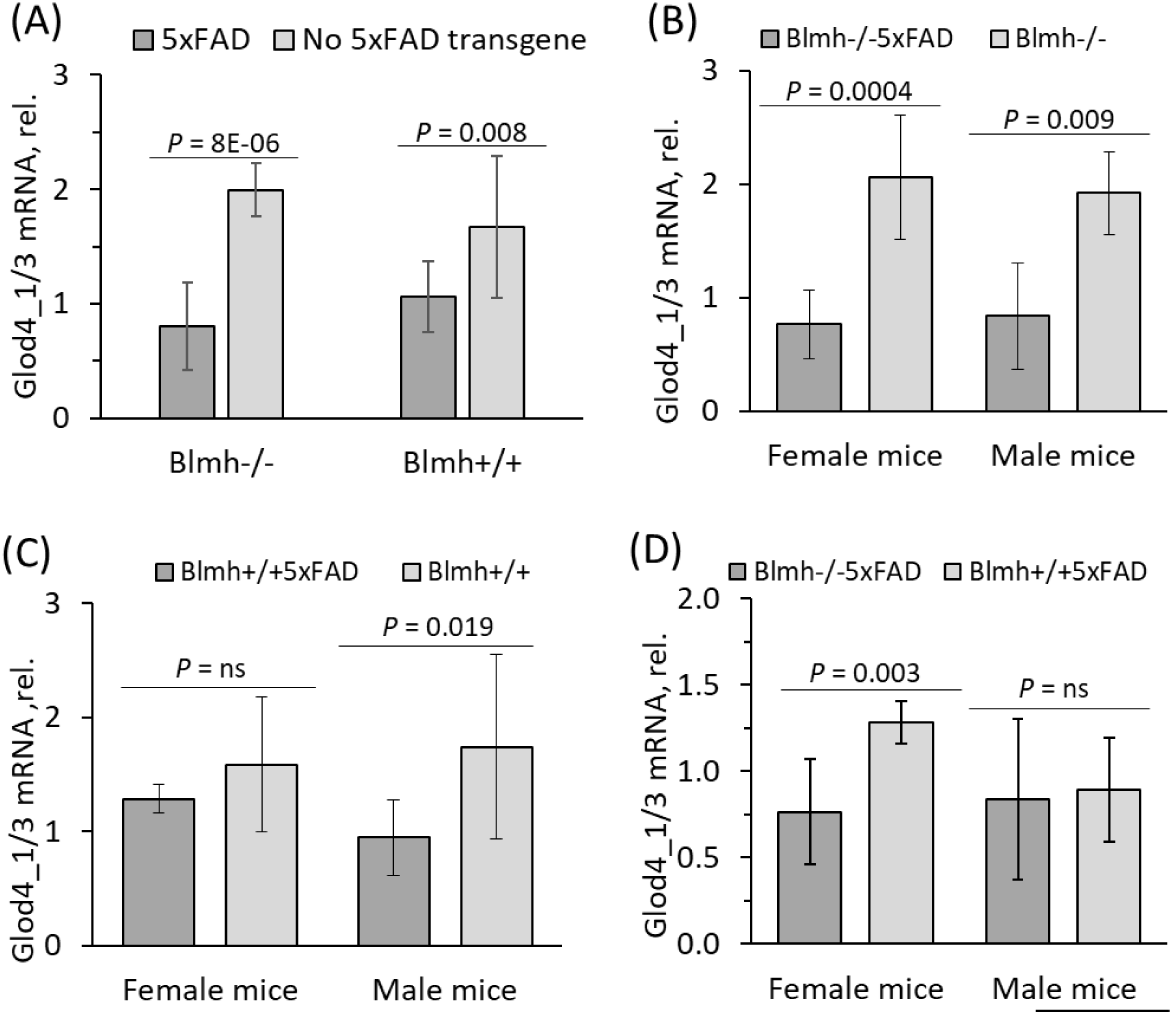
Effects of 5xFAD transgene and sex on *Glod4*_1/3 expression in *Blmh*^−/−^ and wild type *Blmh*^+/+^ mice. Five-month-old *Blmh*^−/−^5xFAD (n = 22), *Blmh*^+/+^5XFAD (n = 15), *Blmh*^−/−^ (n=7), and wild type *Blmh*^+/+^ (n = 4) mice of both sexes were used in the experiments. Glod4_1/3 mRNA was quantified by RT-qPCR using *Actb* mRNA as a reference. (A) 5xFAD transgene reduced *Glod4_1/3* mRNA expression both in *Blmh*^−/−^ and wild type *Blmh*^+/+^ mice. (B) In *Blmh*^−/−^ mice, 5xFAD transgene reduced *Glod4_1/3* mRNA expression independently of sex. (C) In *Blmh*^+/+^ mice, 5xFAD transgene reduced *Glod4_1/3* mRNA expression only in males but not females. (D) *Blmh*^−/−^ genotype reduced *Glod4_1/3* mRNA expression only in 5xFAD females but not males. *p*-values were calculated by an unpaired two-sided Student t test.

*Glod4_1/3* mRNA was significantly downregulated also in *Blmh*^+/+^5xFAD mice compared with non-transgenic wild type *Blmh*^+/+^ animals (**Figure 3A**), although the difference was attenuated compared to the difference seen between *Blmh*^−/−^5xFAD and *Blmh*^−/−^ mice. In contrast to *Blmh*^−/−^ mice, the influence of the 5xFAD transgene on *Glod4* mRNA expression in *Blmh*^+/+^ mice was sex-dependent: observed in males but not in females (**Figure 3C**).

Reduced *Glod4_1/3* mRNA expression, seen in *Blmh*^−/−^5xFAD mice compared to *Blmh*^+/+^5xFAD animals ((0.80 ± 0.38 (n = 22) vs. 1.06 ± 0.31 (n = 15), *P* = 0.045) **Figure 2A, 3A**), was sex-dependent, observed in female but not male mice (**Figure 3D**). However, there was no significant difference in *Glod4_1/3* mRNA expression between six to seven-week-old (**Table 2**) and 5-month-old(1.99 ± 0.23 (n = 4) vs. 1.67±0.62 (n = 7), *P* = 0.405; **Figure 3A**) *Blmh*^−/−^ and wild type *Blmh*^+/+^ mice without the 5xFAD transgene, showing that *Blmh* genotype does not affect Glod4 expression in the absence of the 5xFAD transgene.

### Glod4 gene silencing affects APP and autophagy related Atg5 and p62 mRNAs in mouse neural cells

To elucidate the mechanism by which Glod4 can affect behavioral hallmarks of AD, we first examined whether our findings of downregulated *GLOD4* expression in brains of AD patients (**Figure 1A**) can be recapitulated in mouse neuroblastoma cells N2a-APPswe carrying a mutated human AβPP transgene. We silenced the *Glod4* gene by RNA interference using siRNA Glod4#1 and siRNA Glod4#2 for transfections; controls were transfected with scrambled siRNA (siRNAscr). Gene expression was examined by RT-qPCR using *Actb* mRNA as a reference.

We found that the *Glod4* mRNA level was significantly reduced in *Glod4*-silenced cells (by 76% for siRNA Glod4#1 and by 98% for siRNA Glod4#2; **Figure 4A**). Glod4 protein was also significantly downregulated in the silenced cells (**Figure 4B**). The downregulation of *Glod4* expression was accompanied by upregulation of *AβPP* mRNA (**Figure 4C**) and AβPP protein (**Figure 4D**). Representative images of western blots are shown in **Figure 4E**.

**Figure 4.**
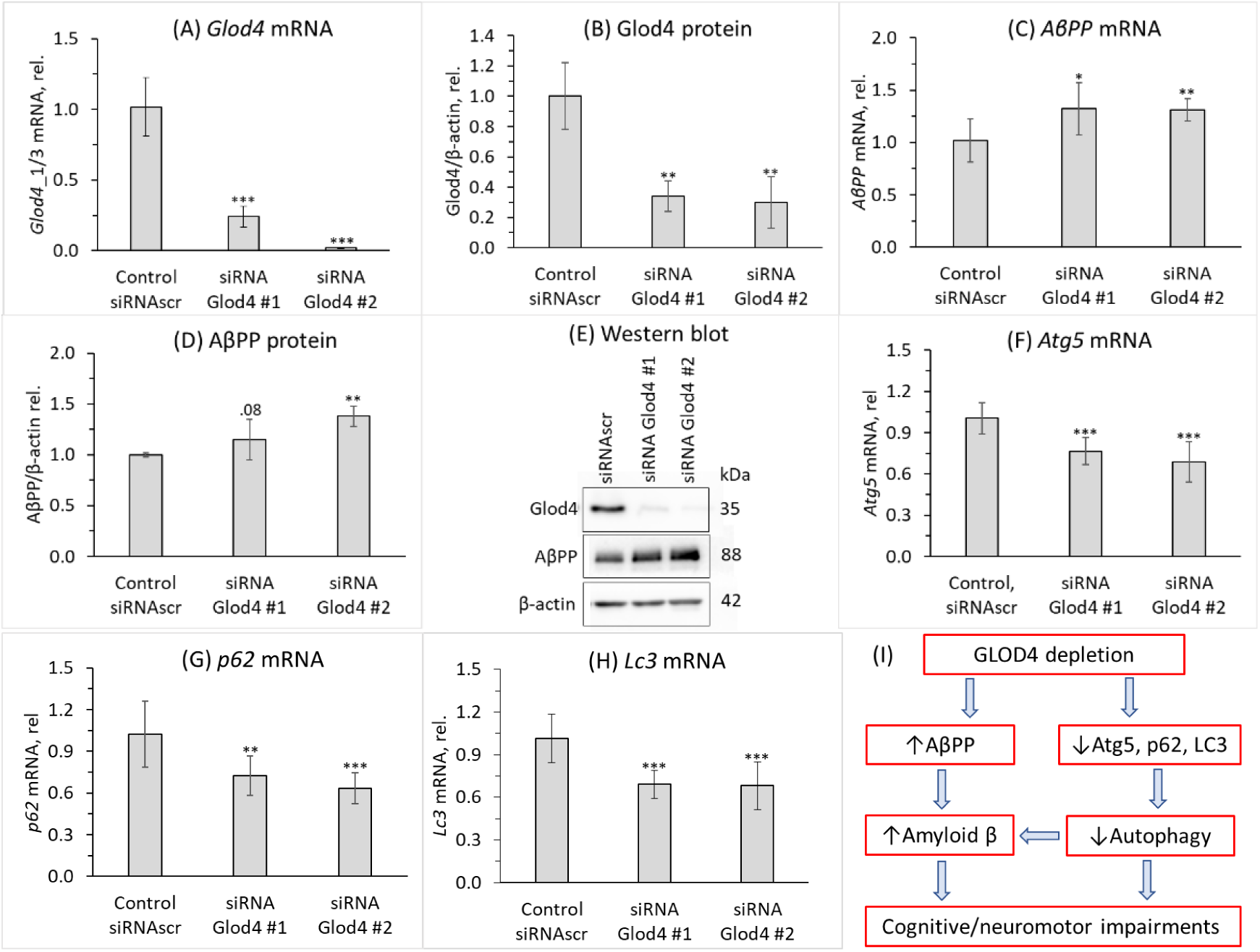
*Glod4* gene silencing affects expression of AβPP and autophagy-related proteins in mouse neuroblastoma N2a-APPswe cells. N2a-APPswe cells were transfected with two different siRNAs targeting the *Glod4* gene (siRNA Glod4 #1 and siRNA Glod4 #2) or scrambled siRNA (siRNAscr) as a control. Proteins and mRNAs were quantified by western blotting and RT-qPCR, respectively, using β-actin protein and mRNA as references. Bar graphs illustrate the quantification of Glod4 mRNA (A), Glod4 protein (B), *AβPP* mRNA (C), and AβPP protein (D). Representative images of western blots are shown in panel (E). Also shown is the quantification of autophagy related mRNAs for *Atg5* (F), *p62* (G), and *LC*3 (H). Each data point is an average ± SD of three technical repeats for each of the three independent biological replicates. Panel (I) illustrates hypothetical pathways by which GLOD4 depletion can contribute to Aβ accumulation and neurological impairments in AD. *p*-values were calculated by an unpaired two-sided Student t test. \**p*<0.05, \*\**p*<0.01, \*\*\**p*<0.001.

Autophagy-related mRNAs for *Atg5* (the regulator of autophagosome assembly) (**Figure 4F**) and *p62* (a receptor for degradation of ubiquitinated substrates) (**Figure 4G**), *Lc3* (microtubule associated protein 1A/1B-light chain 3, involved in formation of autophagosomes) (**Figure 4H**) were significantly downregulated. Other autophagy-related mRNAs such as *Atg7* and *Becn1* were not affected by Glod4 silencing (not shown).

To find whether the effects of Glod4 silencing on autophagy could be mediated by mTOR signaling, we quantified mRNAs levels for *mTOR* and *PHF8* (a positive regulator of mTOR expression) in *Glod4*-silenced and control cells. We found that *mTOR* and *Phf8* mRNA levels were not affected by the *Glod4* gene silencing (not shown).

## DISCUSSION

Our findings suggest that Glod4, a protein with an unknown function, may have a protective role in the CNS. Specifically, we showed that (*i*) *GLOD4* mRNA and protein were downregulated in human AD patients compared to non-AD controls; (*ii*) *Glod4* mRNA was similarly downregulated in *Blmh*^−/−^5xFAD mouse model of AD, mostly in females; (*iii*) The 5xFAD transgene downregulated *Glod4* in *Blmh*^−/−^ mice of both sexes and in *Blmh*^+/+^ males but not females; (i*v*) reduced *Glod4* expression was associated with worsened memory/sensorimotor performance in *Blmh*^−/−^5xFAD mice (*v*); *Glod4* silencing in N2a-APPswe cells upregulated AβPP and downregulated autophagy-related *Atg5*, *p62*, and *LC3* genes. These findings suggest that Aβ accumulation in GLOD4-depleted brains can occur via two pathways shown in **Figure 4I**.

In the first pathway, GLOD4 depletion upregulates AβPP, which increases Aβ accumulation leading to cognitive/neuromotor impairments. In the second pathway, GLOD4 depletion downregulates Atg5, p62, and LC3, which impairs autophagy flux and thus increases Aβ level and cognitive/neuromotor deficits. That impaired autophagy can increase Aβ accumulation mediated by AβPP is known. For example, Becn1, a protein initiating autophagy, is decreased in human AD brains while genetic reduction of Becn1 in transgenic mice that overexpress AβPP (*APP*^+^*Becn*^+/-^ mice) increased Aβ in neuronal cells [28]. However, as we did not find any effect of Glod4 depletion on Becn1 expression (not shown), Glod4 depletion upregulates AβPP and Aβ by a Becn1-independent mechanism.

AβPP level is an important determinant of AD, most strikingly manifested in individuals with Down syndrome who show an early AD neuropathology caused by an extra copy of the *APP* gene due to trisomy of chromosome 21 [29–31]. Transcriptional factors (SP-1, AP-1, CTCF, HSF1,NF-kB, USF, ApoE, androgen) and translational regulators (iron, IL-1, TGFβ, FMRP, hnRNP C, Rck/54, PSF/SFPQ) of AβPP expression have been identified [32]. In previous work, we have shown that deletion of genes encoding enzymes participating in homocysteine metabolism in mice such as Cbs, Pon1, or Blmh elevated levels of AβPP and Aβ in the mouse brain. In the present study, we found that depletion of a protein of unknown function, Glod4, in neural cells (**Figure 4**) and *Blmh*^−/−^5xFAD mice (**Figure 1**) was associated with upregulation of AβPP. These findings identify reduced level of Glod4 as a new factor that can influence the development of AD by upregulating AβPP and Aβ. The mechanism by which Glod4 can regulate AβPP and Aβ levels remains to be elucidated in future studies.

Our findings that Glod4 was depleted in *Blmh^−/−^*5xFAD mice vs. *Blmh*^+/+^5xFAD sibling controls but not in *Blmh*^−/−^ vs. sibling control wild type *Blmh*^+/+^ mice (**Figure 3A**) suggest that Glod4 depletion may be secondary to Aβ accumulation in *Blmh^−/−^*5xFAD animals. However, to find out whether downregulation of Glod4 precedes or is secondary to Aβ accumulation, it would be necessary to examine Aβ and Glod4 levels in *Blmh^−/−^*5xFAD animals at different time points. Our findings also suggest that Hcy-thiolactone and/or *N*-Hcy-proteins, which accumulate in Blmh-deficient mice [33] are unlikely to influence Glod4 expression. The mechanism of Glod4 depletion in mouse *Blmh^−/−^*5xFAD brain and in human AD brain remains to be studied.

Although Blmh depletion caused Glod4 depletion in *Blmh^−/−^*5xFAD mouse brain, downstream effects of Glod4 depletion leading to Aβ accumulation are different from those of Blmh depletion.

Specifically, Blmh depletion in neural cells upregulated mTOR *via* Phf8/H4K20me1 and impaired autophagy by reducing the expression of Atg5, Atlg7, Becn1 and increasing p62 [20]. In contrast, Glod4 depletion did not affect mTOR, Atg7, and Becn1 expression, but downregulated p62 expression (**Figure 4**). Only Atg5 was similarly downregulated in Glod4 and Blmh depleted neural cells. These findings suggest that Glod4 depletion dysregulates autophagy via an mTOR independent mechanism, which remains to be examined.

We found that major GLOD4_1 isoform, and minor GLOD4_2 and GLOD4_3 isoforms, were expressed in the human brain. We also found that all three isoforms were similarly attenuated in AD brains (**Figure 1**). In mice, Glod4_2 isoform was expressed at an extremely low level in wild type animals (0.0008 of the Glod4_1 expression level (**Table 2**) and that Glod4_1/3 isoforms were also downregulated in the *Blmh*^−/−^5xFAD mouse model (**Figure 2** and **3**). These findings suggest that GLOD4 isoforms are unlikely to have an isoform-specific function in AD and thus leave open a question of what the role of GLOD4 isoforms in brain physiology is.

In conclusion, our findings suggest that Glod4 may interact with AβPP and the autophagy pathway, and that disruption of these interactions upregulates Aβ, which causes cognitive/neurosensory deficits, thereby highlighting Glod4’s role in brain homeostasis. Further studies are needed to assess causality of the association of GLOD4 depletion with AD.

## ACKNOWLEDGMENTS

We thank H.G. Lee for samples of AD and control human brains, S.S. Sisodia for the mouse neuroblastoma N2a-APPswe cells, and J. Lazo for the *Blmh*^−/−^ mouse.

## DATA AVAILABILITY

The data that support the findings of this study are available in the methods and/or supplementary material of this article.

